# Probing Patterning in Microbial Consortia with picCASO: a Cellular Automaton for Spatial Organisation

**DOI:** 10.1101/2021.02.14.431138

**Authors:** Sankalpa Venkatraghavan, Sathvik Anantakrishnan, Karthik Raman

## Abstract

Microbial consortia exhibit spatial patterning across diverse environments. Since probing the self-organization of natural microbial communities is limited by their inherent complexity, synthetic models have emerged as attractive alternatives. In this study, we develop novel frameworks of bacterial communication and explore the emergent spatiotemporal organization of microbes. Specifically, we build quorum sensing-mediated models of microbial growth that are utilized to characterize the dynamics of communities from arbitrary initial configurations and establish the effectiveness of our communication strategies in coupling the growth rates of microbes. Our simulations indicate that the behavior of quorum sensing-coupled consortia can be most effectively modulated by the rates of secretion of AHLs. Such a mechanism of control enables the construction of desired relative populations of constituent species in spatially organized populations. Our models accurately recapitulate previous experiments that have investigated pattern formation in synthetic multi-cellular systems. Additionally, our software tool enables the easy implementation and analysis of our frameworks for a variety of initial configurations and simplifies the development of sophisticated gene circuits facilitating distributed computing. Overall, we demonstrate the potential of spatial organization as a tunable parameter in synthetic biology by introducing a communication paradigm based on the location and strength of coupling of microbial strains.

**Author Summary:** Interacting microorganisms that coexist in a given environment tend to have well-defined spatial arrangements. While the emergence of such organization is seen across different microbiomes in nature, it is hitherto not well understood. Decoding the inherent spatial patterning of microbes is constrained by the complexity of their natural habitats. Here, we take advantage of synthetic models of microbial communities to study the dynamics of emergent spatial organization. Our framework of bacterial communication utilizes modular synthetic devices to couple growth rates. In addition to uncovering potential principles of spatial organization, this work enables the construction of complex genetic circuits distributed across communicating strains. Additionally, we have developed a software tool, picCASO, that streamlines the investigation of microbial systems communicating through such frameworks.

## 1 Introduction

Natural microbial communities exhibit spatial organization in numerous niches. Examples include methanogens in mesophilic and thermophilic sludge granules (1), archaea in methane seep sediment (2), nitrifying bacteria in soil aggregates (3), kelp microbiota (4), and the plant rhizosphere (5). In addition, spatial patterning is observed in the human gut, oral, and skin microbiomes, where it is hypothesized to have functional consequences (6–8). The study of organization in these natural microbial communities is limited by the inherent complexity of the system and variable physicochemical conditions. This has driven the design of synthetic communities (9,10) and other *ex vivo* methods (7) to investigate patterning.

Advances in the field of synthetic biology have led to the development of functional systems composed of connected modular elements (11). A key feature of these systems is the interaction between modules that can be intracellular or intercellular. For example, these interactions are intracellular in gene circuits programmed to induce apoptosis in cancer cells (12) and intercellular in engineered mutualistic yeast strains (13). A significant challenge in building complex genetic circuits is the cross-talk that arises while reusing different circuit components within a cell. The distribution of genetic circuits across cells is a strategy used to tackle this challenge (14). However, such compartmentalization across cells requires engineering targeted channels of communication between the cells. In such forms of distributed computing, cellular communication could be optimized by employing spatial organization as a tunable parameter (15).

In this study, we introduce novel frameworks of bacterial communication and explore the emergent spatiotemporal dynamics of microbial consortia. Numerous mathematical models have been developed to study growth in two-dimensional microbial communities, as reviewed previously (16). To explicitly account for spatial heterogeneity, we employ a cellular automaton scheme that has previously been used to model single-species biofilm growth (17), multispecies biofilm growth (18), fungal branching (19), bacterial colony morphologies (20), yeast colony morphologies (21) and colonization of surfaces by bacteria (22).

Our communication paradigms serve as powerful tools to probe the self-organization of microbes. These versatile paradigms, implemented in the form of cellular automaton models, are employed to characterize the growth dynamics of microbial communities from arbitrary initial configurations. We established the biophysical validity of our models by recapitulating experimental data and identified optimal parameters to tune spatial organization through numerical simulations that have shed light on the parameter space. Benchmarking our communication frameworks against well-established models has highlighted their effectiveness in coupling bacterial growth rates. Together, these results provide a solid theoretical basis to understand spatial patterning in microbial consortia.

## 2 Results

Our key results are five-fold. First, we developed a quorum-sensing mediated communication framework for synthetic microbial consortia. Second, we employed a diffusion scheme that accurately reproduced programmed pattern formation in synthetic multicellular systems. Third, we tested the functionality of the model over a wide range of experimentally tractable parameter values using an *in silico* preferential growth assay. Next, we identified a key parameter to modulate patterning in microbial consortia. Finally, we quantified the performance of our quorum sensing-mediated communication strategy and compared its performance to that of a commonly employed metabolite-mediated communication strategy.

### 2.1 Design of Quorum Sensing-Mediated Models of Communication

We developed stochastic agent-based cellular automaton models to investigate principles that govern the spatial organization of microbial consortia. Synthetic quorum-sensing systems with modular components facilitate the coupling of gene expression in any two organisms (23). Our models of quorum sensing-mediated communication were characterized using synthetic two-strain and three-strain *E. coli* communities. Genetically modified *E. coli* strains were ideal for our simulations as *E. coli* is a well-established model organism with high-quality GSMMs and an experimentally validated library of modular quorum-sensing tools. The framework thus established is extensible to all microbes, enabling the study of consortia comprising multiple species.

The quorum sensing-mediated models (QSMMs) comprise coupled *E. coli* strains that constitutively synthesize and secrete acyl-homoserine lactones (AHLs). As depicted in **Fig. 1A**, AHLs secreted by each strain activate gene expression in the coupled strain by inducing a complementary quorum promoter. The strains only differ in terms of the AHLs they produce and the genes driven by the quorum-sensing promoter. Each strain experiences a fitness advantage proportional to its proximity to its coupled strain resulting from the induction of genes downstream of the quorum promoter. Gene expression due to quorum sensing is considered to be a Boolean process gated by an AHL concentration threshold, consistent with experimental data (23). This allows for the coupling of multiple strains in a microbial consortium through quorum sensing-driven communication and gene expression. An orthogonal coupling paradigm is employed in our multistrain models (**Fig. 1B**).

**Figure 1:**
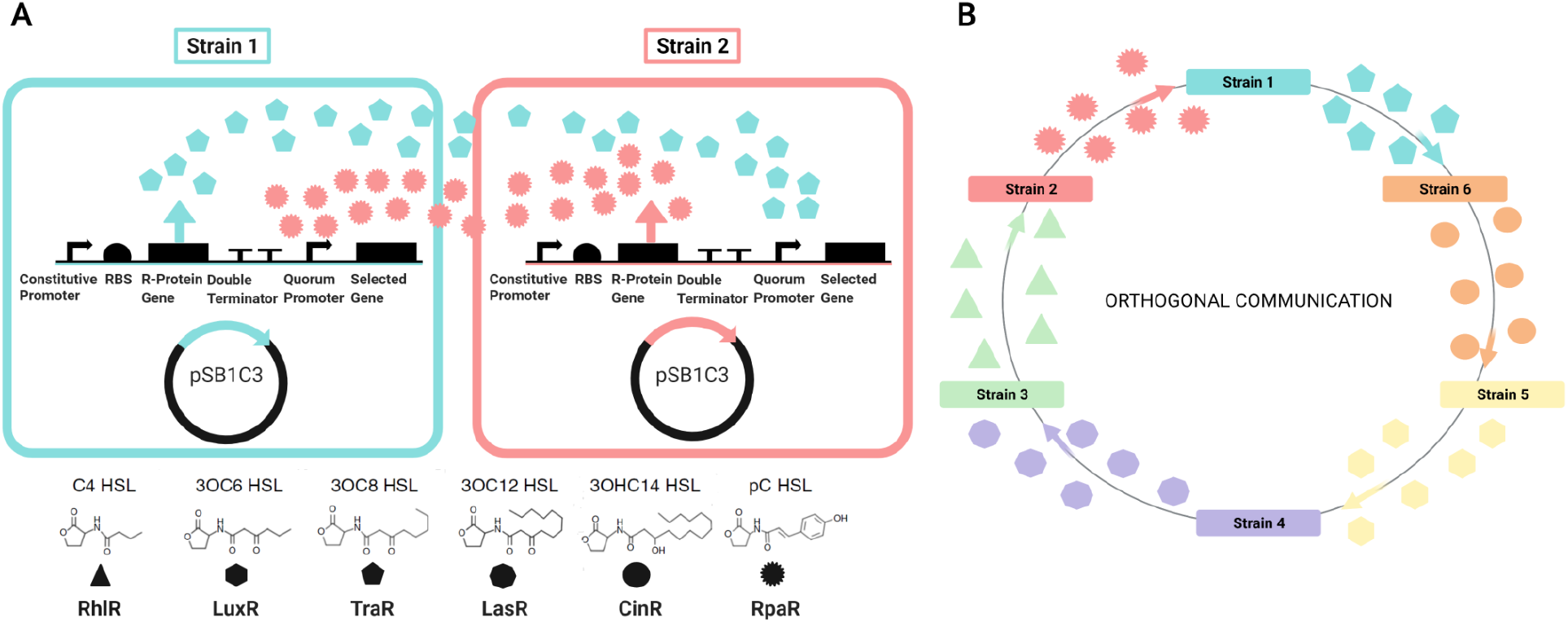
Schematic representation of the genetic circuit designed to implement the quorum sensing mediated model (QSMM) for coupling bacterial strains using acetyl-homoserine lactones (AHLs) and quorum sensing module pairs. (A) Strain 1 secretes 30C8 HSL that causes gene expression in Strain 2 by activating TraR. Strain 2 secretes pC HSL which in turn causes gene expression in Strain 1 by activating RpaR. (B) Orthogonal coupling between six strains, indicating how this paradigm can be extended to study any microbial community of interest. AHLs C4 HSL, 3OC6 HSL, 3OC8 HSL, 3OC12 HSL, 3OHC14 HSL, and pC HSL are represented by triangles, hexagons, pentagons, octagons, circles, and spiked circles respectively. Figure created using BioRender.

Our FBA-based strategy quantifies the contribution of genes driven by the quorum promoter to the fitness of a strain—defined in terms of its growth rate (**Methods**). This notion is formalized by measuring the reduction in growth rate with respect to the wild-type strain upon the deletion of each gene from a GSMM. We performed single-gene deletions in *E. coli* iAF1260 and identified 27 non-lethal genes that resulted in reduced growth rates ranging from 30–90% (**S2 Table**). The genetic backgrounds of strains in our study are such single-gene knockouts with reduced growth rates. Quorum sensing results in the re-expression of the knocked-out gene and rescues the wild-type growth rate. Therefore, a strain is expected to have a higher growth rate in the presence of its coupled strain.

We utilize the FBA-based strategy to define strains in our two-strain QSMM consisting of strain 1 (blue) and strain 2 (red), and a three-strain QSMM comprising strain 1 (blue), strain 2 (red), and strain 3 (green). An orthogonal coupling strategy is employed in the three-strain model in which AHLs secreted by strain 1 are sensed by strain 3, those secreted by strain 2 are taken up by strain 1, and those secreted by strain 3 induce gene expression in strain 2. The initial values of all parameters in these strains are taken to be the same.

### 2.2 Model Accurately Reproduces Patterning in Synthetic Multi-cellular Systems

Our models of microbial growth were simulated on two-dimensional grids. We defined cell-blocks to be squares on the grid containing approximately 10^8^ *E. coli* cells in an area of 10 mm^2^ (24). Each cell secretes AHLs at a rate of 10^-8^ nmol/h (25). We modelled the secretion and subsequent distribution of AHLs in this grid using a Forward-Time Central Space (FTCS) scheme of diffusion. Each time-step in our simulations is 10 minutes (**Methods**). In every step, an amount of AHL proportional to the amounts present in neighboring squares enters a given square in the grid. Consequently, AHL effluxes proportional to the amount present in a given square takes place to each of its neighbors. The diffusivity of AHL is taken to be 8×10^-10^ m^2^/s (26). The decay of AHL molecules follows first-order kinetics with a rate constant of approximately 4.6×10^-4^ min^-1^ (27). Similar schemes have been employed to model diffusion in two-dimensional microbial cultures (21).

We validated our model definition and diffusion scheme by replicating results from Basu *et al*. (2005). Basu *et al*. developed a band-pass filter in which fluorescent proteins are only expressed within a narrow AHL concentration range (0.01–0.2 μM). Their quantitative experiments were performed on a solid medium containing a lawn of “receiver” bacteria sensing AHLs emitted by a “sender” colony seeded in the middle. In approximately 36 hours, a steady-state was reached in which cells fluoresced in a ring between 5 and 18 mm from the sender colony. We translated these experiments in the context of our model using a grid with a central “sender” cell-block surrounded by a lawn of “receiver” cell-blocks. Only receiver cell-blocks exposed to AHL concentrations within the band-pass filter window fluoresce (**Fig. 2A**). The ring of fluorescence was tracked over time until a stable pattern was observed.

**Figure 2:**
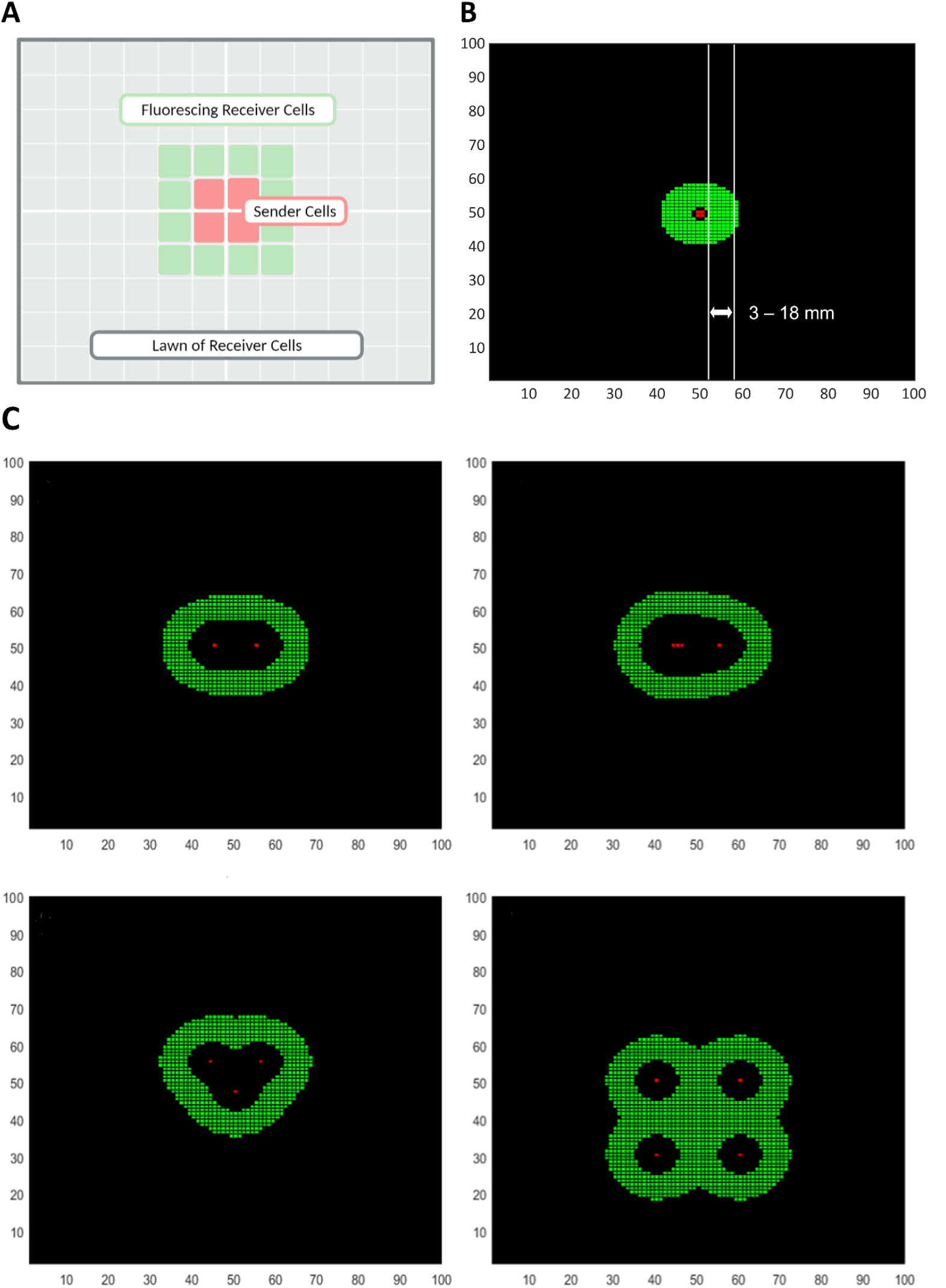
Validation of the FTCS diffusion scheme for quorum sensing molecules. (A) A schematic representation of the simulation setup, with four sender cell-blocks secretingAHLs amidstalawn ofreceiver cell-blocks that fluoresce when the local AHL concentration falls within a defined range (0.01–0.2 μM). (B) Simulation results capturing the quantitative characteristics ofthe GFP bandpass filter described in Basu et al. (2005) resulting in a band (shown in green) ranging from 3–18 mm from the sender colony (shown in red). (C) Simulation results qualitatively reproducing four patterns observed by Basu *et al*. (2005) by seeding sender colonies of different sizes in different configurations on a lawn of receiver cells.

As seen in **Fig. 2B**, fluorescent gene expression was observed in a band spanning 3–18 mm (2–6 blocks) from the sender colony. The resolution of our coarse-grained model is 3 mm, which explains the minor discrepancy in the inner radius of the fluorescent ring. This stable pattern was achieved after 216 time-steps, corresponding to a period of 36 hours, recapitulating experimental data (28). Additionally, we recreated several other patterns obtained from the’ qualitative experiments of Basu *et al*. by seeding sender colonies in different positions across the grid (Fig 2C, Fig. 5 from Basu *et al*. (2005)). These results substantiate the values of model parameters obtained from literature including the density of cell-blocks, the time-step, the rate of secretion of AHLs, the rate of degradation of AHLs, and the diffusivity of AHLs in agar gels.

**Figure 3:**
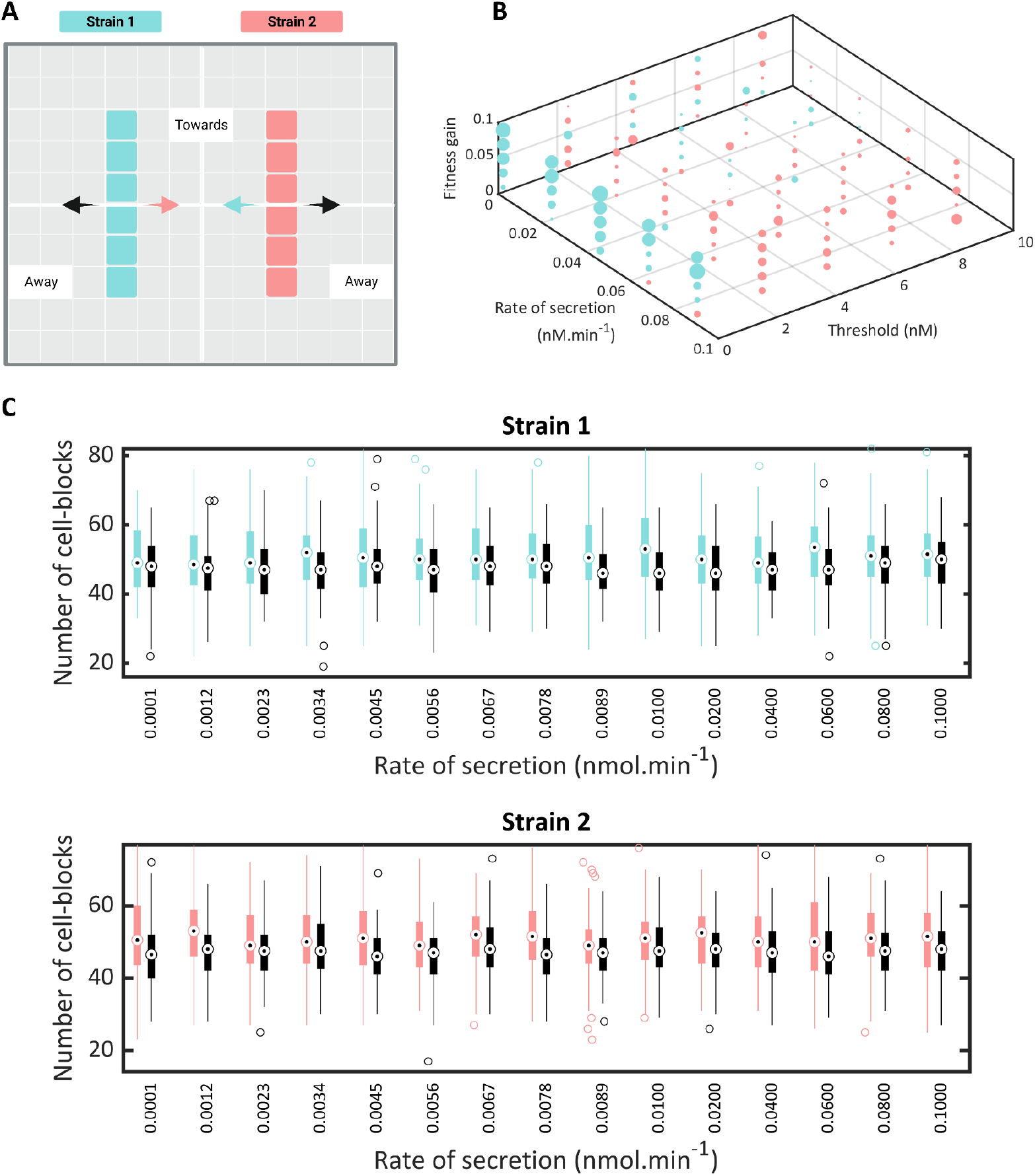
A grid search and preferential growth assays reveal model behavior and stability across parameter space. (A) A schematic representation of the simulation setup for the preferential growth assay with cell-blocks of two coupled strains seeded in parallel lines. Coupling between the strains is expected to cause cell-blocks from one strain to proliferate toward the other. (B) A 3D plot depicting the variation in the number of cell-blocks of a strain with respect to Strain 1’s rate of secretion of QS molecules, the gain in fitness, and the threshold concentration required to induce gene expression in the QSMM model with two strains. The color of each point in the plot corresponds to the strain with the higher number of cell-blocks, following the same color scheme as in (A), while its size is proportional to the difference between the number of cell-blocks of each strain. (C) Results of a preferential growth assay quantifying the number of cell-blocks of Strain 1 (above) and Strain 2 (below) dividing towards (shown in color) and away from (shown in black) cell-blocks of the other strain with the rate of secretion of AHL molecules from Strain 1 being varied and the rate of secretion from Strain 2 kept constant at 1.6×10^-2^ nM cell-block^-1^ min^-1^.

**Figure 4:**
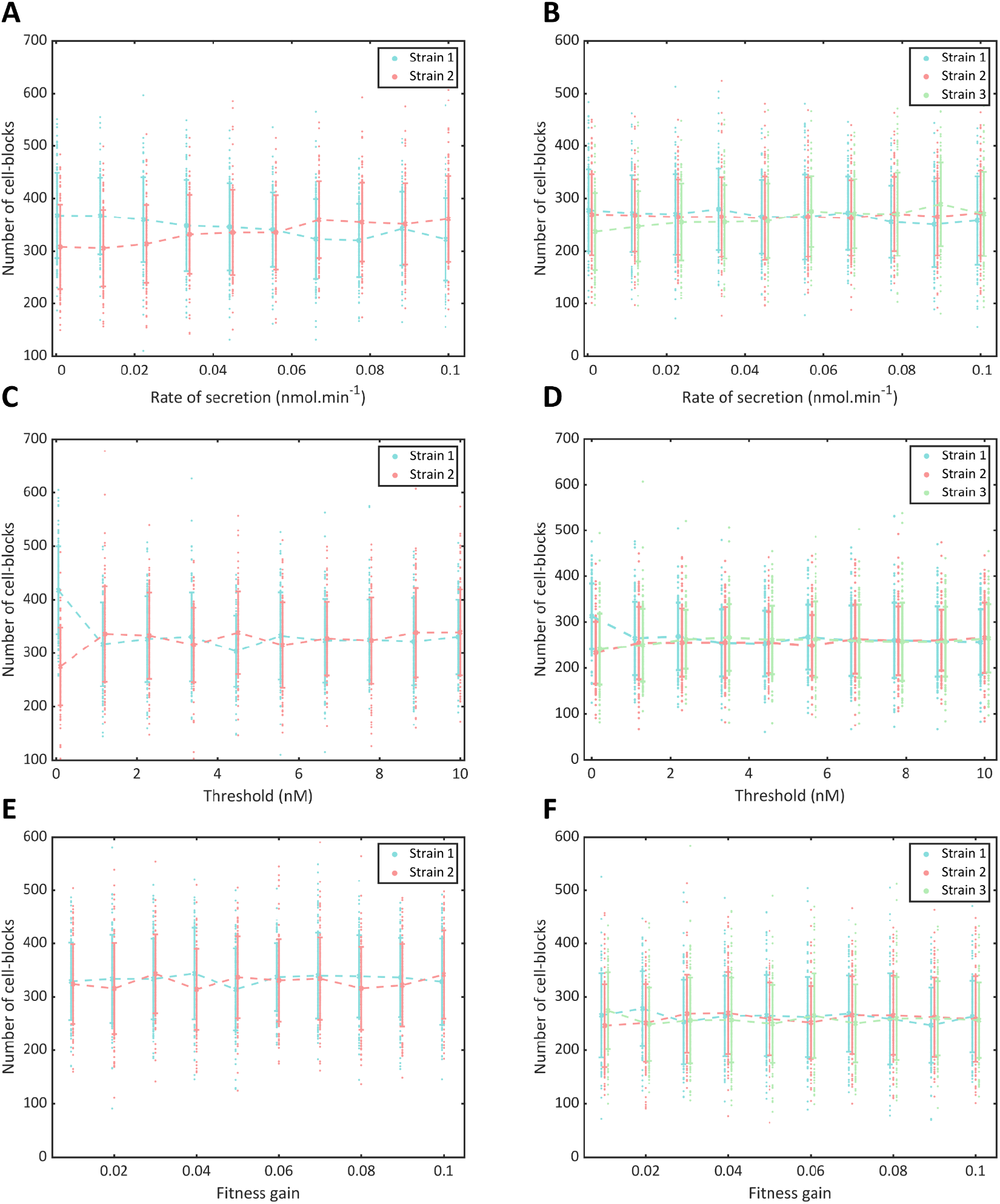
Variation in model behavior obtained by varying a single parameter. (A, B) The populations of each strain in the (A) two-strain and (B) three-strain QSMM with the rate of secretion of AHL molecules from Strain 1 being varied and the rate of secretion from the other strain(s) kept constant at 1.6×10^-2^ nM cell-block^-1^ min^-1^. (C, D) The populations of each strain in the (C) two-strain and (D) three-strain QSMM with the threshold AHL amount to induce gene expression in Strain 1 being varied and the threshold for the other strain(s) kept constant at 5 nM. (E, F) The populations of each strain in the (E) two-strain and (F) three-strain QSMM with the fitness gain due to QS-mediated gene expression in Strain 1 being varied and the fitness gain for the other strain(s) kept constant at 0.05.

**Figure 5:**
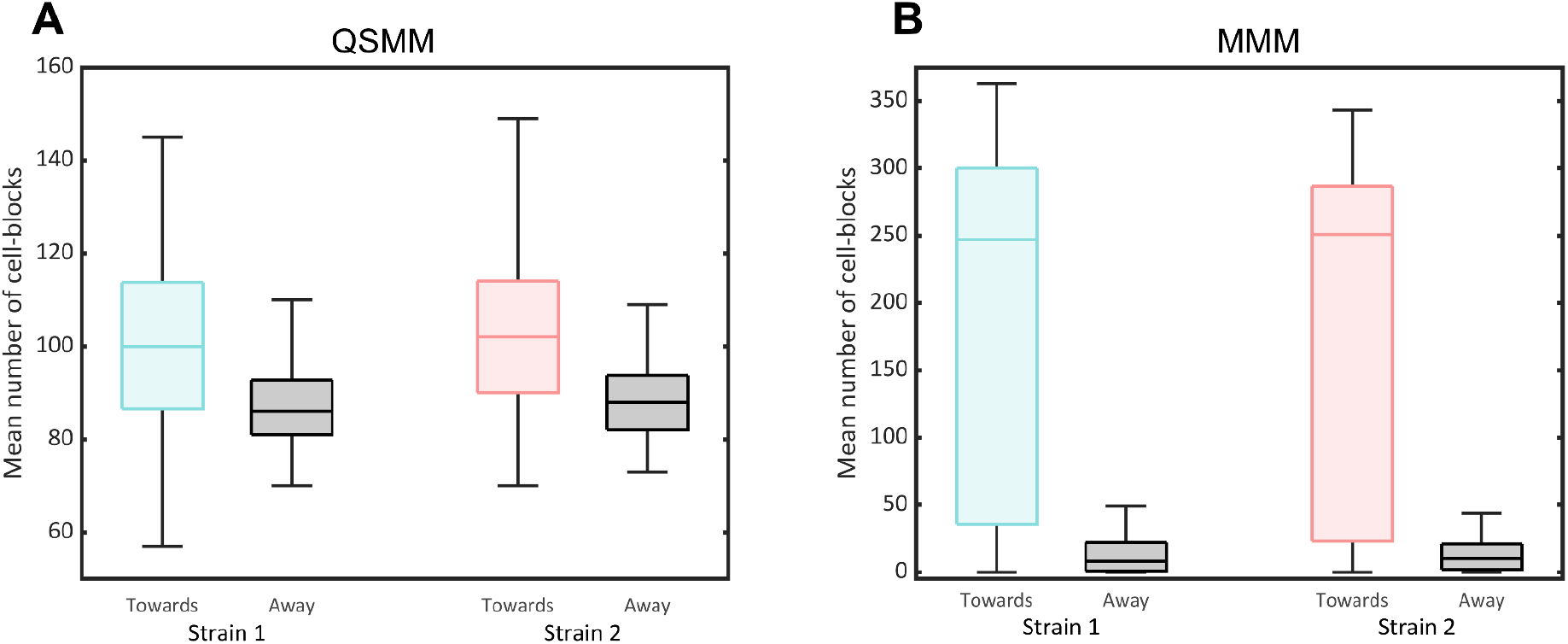
Comparison of the strength of coupling. The strengths of coupling are determined in (A) the QSMM and (B) the MMM through quantification of the number of cell-blocks of a given strain that divide towards and away from the second strain in a two-strain model using the preferential growth assay. For the QSMM, *p* = 10^-11^ for Strain 1 and *p* = 10^-12^ for Strain 2. For the MMM, *p* = 10^-24^ for Strain 1 and *p* = 10^-23^ for Strain 2.

### 2.3 Observation of Coupling Between Strains Over a Wide Range of Experimentally Tractable Parameters

Next, we sought to establish that the strains in our models were coupled through quorum sensing-mediated communication. We designed a preferential growth assay based on the principle that strains have a higher growth rate proximal to their coupled strain. In this assay, cell-blocks consisting of a pair of coupled strains were seeded in parallel lines on a grid (**Fig. 3A**). The rates of proliferation of each strain towards and away from the coupled strain were measured. In the absence of coupling, it is expected that these rates will be nearly equal while the presence of coupling would increase the rate of proliferation of a strain towards its coupled strain.

The preferential growth assay was not only applied to our initial model set-up but was also used to test model functionality for experimental applications. To facilitate a systematic analysis, we chose to vary the parameters that are most amenable to experimental modification over a tractable range of values for one strain while maintaining the default values for the coupled strain. The rate of secretion of AHLs from a cell can be modulated by driving the expression of AHL-synthesizing enzymes with promoters of different strengths. This parameter was varied from 10^-12^ to 10^-9^ nM/cell·min with a default value of 1.6×10^-10^ nM/cell·min (25). The threshold AHL concentration to induce gene expression was set between 10^-1^ and 10^4^ nM, with a default value of 5 nM (29). This parameter can be tuned by using different modular quorum sensing systems. The quorum sensing-induced fitness gain can be adjusted by employing our FBA-based strategy to select appropriate strains. The default value of the fitness gain was 0.05, and it was varied over a range from 0.01 to 0.1 (**Methods**).

The number of cell-blocks of each strain that have proliferated towards and away from the coupled strain is visualized in **Fig. 3C**. The mean number of cell-blocks that have proliferated towards the coupled strain is higher for both strain 1 and strain 2 across all parameter values studied. We observed that for all three parameters studied (**Fig. 3C, Supp. Fig. 1, Supp. Fig. 2**), significant preferential growth is seen across the entire range being considered. This demonstrates the presence of significant coupling over all experimentally tractable ranges of parameters.

### 2.4 AHL Secretion Rate Optimally Modulates the Behavior of Quorum Sensing-mediated Consortia

We studied the behavior of coupled QSMM systems in the parameter space after establishing that strains in our models exhibit coupling across the range of experimentally tractable parameter values. For the two-strain model, a grid search was performed in which we varied the values of the rate of secretion of AHLs (10^-4^ to 10^-1^ nM/cell-block·min), the quorum sensing threshold concentration (10^-1^ and 10^4^ nM), and the fitness gain (0.01–0.1) simultaneously. These parameters were varied for strain 1 and kept constant at the aforementioned default values for strain 2. Initially, four cell-blocks of each strain were seeded randomly on a grid and allowed to proliferate for 50 steps, corresponding to 500 minutes. The model was iterated 100 times and the mean numbers of cell-blocks of the two strains at the end of each simulation were computed and visualized in **Fig. 3B** and **Supp. Fig. 3**.

Each point in **Fig. 3B** represents an average of over 100 iterations of the model for a parameter set. The color of each point corresponds to the strain with a higher number of cell-blocks and its size is proportional to the difference between the number of cell-blocks of both strains. We observe that when the rate of secretion from strain 1 increases, the proportion of strain 2 in the population increases. Strain 2 dominates when strain 1’s threshold AHL concentration is very low, but this effect disappears at higher thresholds. Further, the effect of fitness gain was seen to be negligible. Together, these observations indicate that community behavior is most effectively tuned by modulating the rate of secretion. The inherent stochasticity in our models is reflective of the random processes that govern microbial growth and spatial patterning. While model insights are statistically significant, the precise role played by stochasticity remains to be elucidated. We anticipate that the effects of stochasticity will reduce over longer simulation times.

To obtain a more fine-grained picture of the above results, we varied each parameter individually for strain 1 over the same range with 4 cell-blocks initially seeded randomly on the grid and allowed to proliferate for 100 steps (1000 minutes) for 100 runs of the model. These simulations were performed for the two-strain and three-strain models (**Fig. 4**). Consistent with the above results for the two-strain models, we notice that a higher rate of secretion from strain 1 led to an increase in the number of cells of strain 2. Similar trends were observed in the case of the three-strain model. As expected, a higher rate of secretion of strain 1 resulted in an increase in the mean number of cell-blocks of strain 3 and no significant change in strain 2. At very low threshold concentrations for strain 1, the fraction of strain 1 in the population was very high in both the two-strain and three-strain models. However, the populations of all strains rapidly converged to nearly equal values when the threshold was increased. This could be because the threshold values far exceed the AHL concentrations observed in the timeframe of the simulation, resulting in no strain experiencing a fitness advantage. Variation of the fitness gain upon quorum sensing shows no discernible trend, perhaps due to the stochastic nature of the processes involved. Therefore, we observe that among the parameters considered, tuning the rate of secretion of quorum sensing molecules is most advantageous to regulate consortium behavior.

### 2.5 Benchmarking the Quorum Sensing Based Coupling Strategy

We have implemented quorum sensing mediated communication between strains in a consortium, studied model behavior, and derived insights on tunable parameters. Conventionally, communication between microbial strains has been engineered through the creation of auxotrophic strains that exchange essential metabolites (13). To compare our models with other modes of engineering communication between strains we employ a metabolite-mediated model (MMM). In the two-strain MMM, each strain is an auxotroph that is incapable of producing an essential metabolite. Its coupled strain produces and secretes this metabolite, thereby facilitating its growth and proliferation. Thus, these auxotrophic strains cannot grow in the absence of their mutualistic counterparts. The diffusivity of metabolites is taken to be 5×10^-10^ m^2^/s, which is the diffusivity of maltose (30). As this value is representative of the diffusivity of metabolites, our results are extensible to a broader range of common metabolites. For the MMM, the base fitness is set to zero, as a strain cannot proliferate in the absence of an essential metabolite.

To compare the strategies of coupling employed in the QSMM and the MMM, we performed preferential growth assays using two-strain models (**Fig. 3A, Fig. 5**). In both models, the mean number of cell-blocks of each strain proliferating towards its coupled strain was higher than that proliferating away from the coupled strain. This observation was statistically significant for both the QSMM and MMM. The fraction of each strain proliferating towards and away from the coupled strain in the MMM was an order of magnitude higher. Therefore, the strength of coupling was far greater in the case of the MMM. Nevertheless, our results indicate that the strength of coupling in the QSMM is significant and that this method can be used to engineer communication in consortia.

## 3 Discussion

In this study, we have designed a novel quorum sensing-mediated communication framework, modeled interactions in synthetic microbial consortia, and harnessed this strategy to study spatial organization. We devised an FBA-based strategy to enable the coupling of growth rates of designer strains in a microbial community. Our models have facilitated the study of two-dimensional spatial patterning over a wide range of experimentally tractable parameter values. These simulations can be flexibly employed to characterize the growth dynamics of consortia with any arbitrary initial configuration. This work demonstrates the use of spatial organization as a tunable parameter in synthetic biology by engineering communication based on the location and strength of coupling of microbial strains. Engineering organization in synthetic microbial consortia has far-reaching applications. Controlling spatial patterning allows for the compartmentalization of different functional genetic circuits, which is an effective means of reducing both the biochemical crosstalk and the complexity of synthetic genetic circuits within each microbe. A direct consequence of this study is the ability to develop more sophisticated gene circuits for distributed computing. Additionally, our models can be built upon to study the behavior of natural microbial consortia. Therefore, our work enables the investigation of spatial patterning in natural microbial communities and provides a means to study population dynamics in consortia.

Our software tool is generalizable to any species in a consortium through the use of appropriate genome-scale metabolic models, growth rates, and AHL secretion rates. Similarly, diverse environments can be modeled by employing suitable diffusion coefficients and rates of small molecule degradation. A potential drawback of quorum sensing-mediated communication systems is the non-specific activation of downstream targets resulting from cross-talk. We envision the use of our models in conjunction with the computer-aided-design tool developed by Polizzi and co-workers which facilitates the selection of orthogonal AHL systems from an experimentally validated library, resulting in reduced cross-talk (23). A limitation of our models is that they do not account for intercellular dynamics, and thus cannot accurately recapitulate some temporal aspects of gene expression. The use of a cellular automaton scheme is advantageous to study cellular dynamics as compared to differential equation-based models because it explicitly accounts for spatial heterogeneity. Due to the computational infeasibility of simulating the behavior of each cell in agent-based models, we employed a cell-block approximation. Cell-block size is defined to enable doubling within a single time-step, cell-block density is derived from experimental data and the rates of secretion of small molecules are scaled appropriately. Together, these criteria ensure the physical validity of the approximation and reduce the computational complexity of simulating large consortia.

Overall, we show that the behavior of quorum sensing-coupled consortia can be effectively modulated by controlling the rates of secretion of AHLs. This mechanism of control enables the construction of spatially organized populations with desired relative populations of constituent species. We performed a systematic evaluation of the communication strategy employed in the QSMM by benchmarking it against the MMM. Strains were found to be coupled in both models, but the strength of coupling was higher in the MMM. This can be understood intuitively as the MMM uses auxotrophic strains whose base fitness, in the absence of the mutualistic strain, is zero. In the QSMM, on the other hand, strains have a low but non-zero base fitness even in the absence of other strains. An essential component of engineered microbial consortia is the communication strategy employed. QSMM-based communication systems are modular and can be inserted into an organism in a plug-and-play manner. The QSMM offers a large repertoire of AHLs due to the availability of experimentally validated libraries. The MMM is restricted by the number of metabolites that are essential and can be secreted and taken up by the microbes in the consortium.

In summary, we established versatile models, available as an easy-to-use MATLAB package, that enables the study of spatial patterning in microbial communities. They provide a theoretical basis to understand the self-organization of microbes in consortia. A potential extension of our work is the development of tools to calculate initial seed configurations to arrive at specified patterns. Such a tool would provide predictive power to our model and allow experimentalists to generate patterns of interest. To conclude, our microbial communication strategy is scalable and facilitates the design of consortia for applications across distributed circuit design, designer microbiomes, and biofilm manipulation.

## 4 Materials and Methods

### 4.1 Model Framework

We implemented all models in MATLAB (Mathworks Inc., USA) using a two-dimensional grid in which the nearest neighbors are considered as the four non-diagonally-adjacent squares. The replicating units in our models are defined as cell-blocks consisting of 10^8^ cells to simplify computation (21). They are represented using classes that contain methods to identify empty neighboring blocks and simulate cell division. Cell-blocks of strains engineered with quorum sensing modules are seeded onto the grid in numerous user-defined configurations in our simulations. The diffusion of AHLs secreted by each strain is implemented using a Forward-Time Central-Space (FTCS) scheme (31).

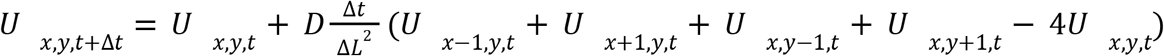

where *U*_*x*,*y*,*t*_ is the concentration of the resource in the square at the coordinates (*x*,*y*) at time *t*, Δ*t* is the time step, Δ*L* is the square length, and *D* is the diffusion constant for the resource. The degradation of these molecules is assumed to follow first-order kinetics.

Each strain is defined to have an intrinsic base fitness that increases upon gene expression due to quorum sensing. Cell division occurs probabilistically depending on the fitness of the cell-block. At each time step, the fitness of each cell-block is re-computed based on the local concentration of relevant AHLs. Thus, cell-blocks with higher fitnesses are more likely to divide. The model has been implemented for consortia consisting of two and three distinct *E. coli* strains but is readily extendable to microbial communities comprising both different species and more strains.

### 4.2 Strain Design

The strains used in our quorum sensing mediated communication framework contain a single gene deletion that results in a reduced growth rate compared to the wild type. Re-expression of a knocked-out gene through quorum sensing restores the wild-type growth rate, which is modeled as an increase in the fitness of the cell-block.

Flux Balance Analysis (FBA) is a technique used to predict the growth rate and metabolic capabilities of cells using genome-scale metabolic models (32,33). Under conditions of steady-state mass balance, FBA predicts the growth rate of cells and possible flux states that correspond to maximum growth. Additionally, FBA can be used to characterize the phenotype of a cell following perturbations such as a gene deletion (34).

We used FBA to compute the growth rates, in the form of biomass production rates, of wild-type cells using Genome-Scale Metabolic Models (GSMMs). Our simulations are performed using the COBRA toolbox for MATLAB (35). Single gene deletions are simulated *in silico* and the growth rates of each knockout strain relative to the wild-type are quantified. Genes that resulted in a reduced growth rate, that is 30–90% of the wild-type growth rate, were shortlisted as candidate genes for further simulations. The reduced growth rates obtained were used to compute the base fitness of each strain. The wild-type fitness is defined as the time step per iteration divided by the generation time. This leads to the cell-block dividing once per generation time on average. The base fitness is taken to be the wild-type fitness multiplied by the growth ratio, i.e., the ratio of the biomass production rate of the knock-out strain to that of the wild-type. The design of strains was facilitated by a library of pairs of AHL - quorum sensing modules with well-characterized threshold concentrations developed by Kylilis *et al*. (2018).

### 4.3 Software Tool

Our models are available as a package at https://github.com/RamanLab/picCASO. The *E. coli* iAF1260 GSMM is the default genetic background of strains (36). Additionally, users have the option of simulating any microbial species by inputting its growth rate and corresponding GSMM. Following this, strains are designed with genes to be knocked out selected using the FBA-based strategy and with appropriate AHL - quorum sensing module pairs selected by the user. The biophysical parameters of the chosen pairs are incorporated into the classes corresponding to the cell-blocks of each strain. Predefined seed cell-block configurations including arrangements in parallel lines, in a single file, at diagonally opposite corners of a square, in concentric circles, and in random distributions, are available for the user to select from. Further, provisions have been made for users to encode custom patterns of interest and analyze their dynamics.

## 5 Acknowledgements

KR acknowledges support from the Science and Engineering Board (SERB) MATRICS Grant MTR/2020/000490, IIT Madras, Centre for Integrative Biology and Systems Medicine (IBSE), and Robert Bosch Center for Data Science and Artificial Intelligence (RBC-DSAI). SA acknowledges The Department of Science and Technology (DST) for the INSPIRE Scholarship for Higher Education (SHE). The authors acknowledge the use of the computing resources at HPCE, IIT Madras.

