## Supplementary Information for "Probing Patterning in Microbial Consortia with picCASO: a Cellular Automaton for Spatial Organisation"

| **Parameter** | **Value** | **Reference** |
|---|---|---|
| Diffusion coefficient (D) - Quorum Sensing Molecules | $8\times10^{-10}$ $m^2/s$ | (1) |
| Diffusion coefficient (D) - Metabolites | $5\times10^{-10}$ $m^2/s$ | (2) |
| Base fitness | User-defined ($10/T_d$ in mins)<br>Default: 0.082 | Methods |
| Threshold | User-defined<br>Default: 1 nM | (3) |
| Fitness changes due to quorum sensing | User-Defined<br>Default: 0.082 | Methods |
| Rates of secretion of quorum sensing molecules (QS_AMT_1, QS_AMT_2, and QS_AMT_3) | 0.16 nmol/cellblock.iter | (4) |
| Rate of degradation of quorum sensing molecules (dgAHL) | 0.0046 per iteration | (5) |
| Distance between neighbouring cells (dl) | $3\times10^{-3}$ m | Methods |
| Time elapsed per iteration (dt) | 600s | Methods |
| Grid size (gridlenx × gridleny) | Default: 100x100 | Methods |
| niter | Default: 100 | Methods |

**S1 Table:** Default values of parameters used in simulations.

| # | Gene ID | Gene Name | grRatio |
|---|---|---|---|
| 66 | 'b3733' | 'atpG' | 0.3220276 |
| 67 | 'b3735' | 'atpH' | 0.3220276 |
| 71 | 'b3731' | 'atpC' | 0.3220276 |
| 72 | 'b3734' | 'atpA' | 0.3220276 |
| 73 | 'b3738' | 'atpB' | 0.3220276 |
| 74 | 'b3732' | 'atpD' | 0.3220276 |
| 75 | 'b3736' | 'atpF' | 0.3220276 |
| 76 | 'b3737' | 'atpE' | 0.3220276 |
| 273 | 'b0429' | 'cyoD' | 0.860351 |
| 275 | 'b0432' | 'cyoA' | 0.860351 |
| 280 | 'b0431' | 'cyoB' | 0.860351 |
| 281 | 'b0430' | 'cyoC' | 0.860351 |
| 386 | 'b1779' | 'gapA' | 0.7880651 |
| 857 | 'b2926' | 'pgk' | 0.7880651 |
| 1024 | 'b2279' | 'nuoK' | 0.8158387 |
| 1026 | 'b2283' | 'nuoG' | 0.8158387 |
| 1028 | 'b2278' | 'nuoL' | 0.8158387 |
| 1032 | 'b2288' | 'nuoA' | 0.8158387 |
| 1034 | 'b2284' | 'nuoF' | 0.8158387 |
| 1036 | 'b2285' | 'nuoE' | 0.8158387 |
| 1040 | 'b2281' | 'nuoI' | 0.8158387 |
| 1042 | 'b2280' | 'nuoJ' | 0.8158387 |
| 1045 | 'b2277' | 'nuoM' | 0.8158387 |
| 1047 | 'b2282' | 'nuoH' | 0.8158387 |

| | | | |
|---|---|---|---|
| 1050 | 'b2286' | 'nuoC' | 0.8158387 |
| 1052 | 'b2276' | 'nuoN' | 0.8158387 |
| 1056 | 'b2287' | 'nuoB' | 0.8158387 |

**S2 Table:** Gene list for *E. coli* iAF1260 obtained from FBA studies. grRatio is the normalised growth rate of the cell with respect to the wildtype, obtained from single gene deletions filtered to select for those in the range 30-90%.

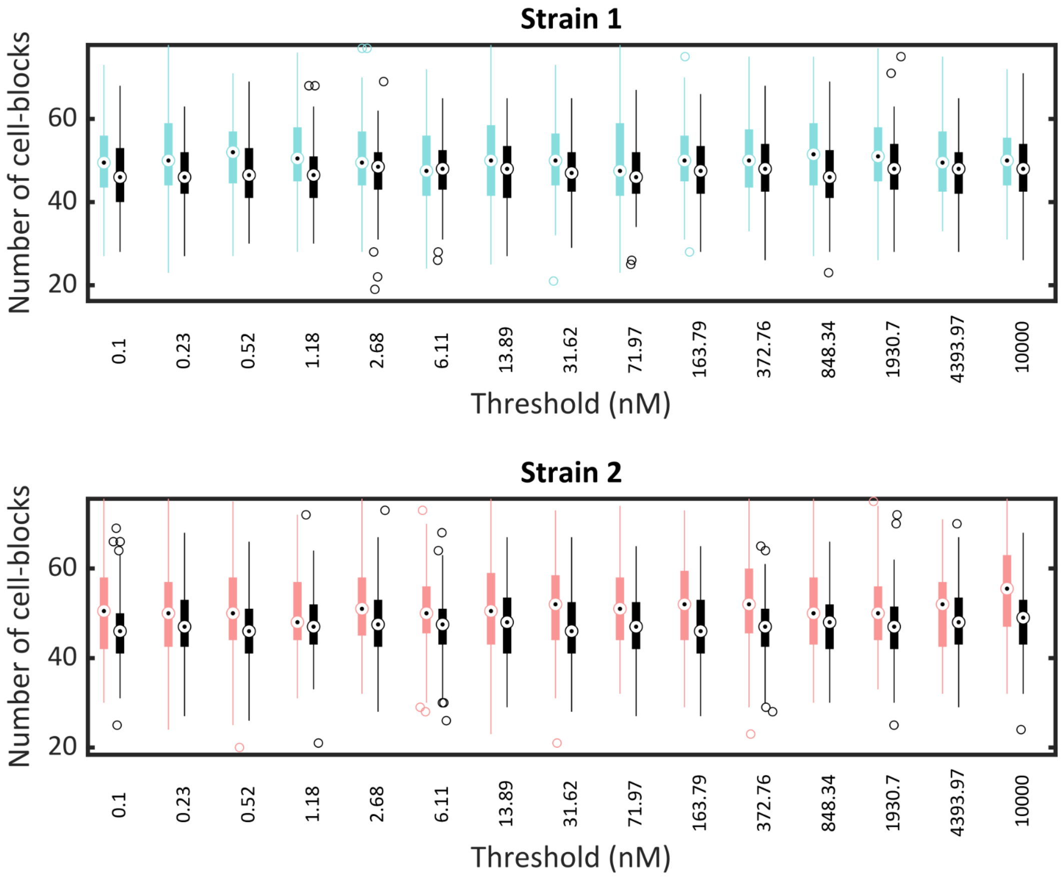


**Supp. Fig. 1**: Results of a preferential growth assay quantifying the number of cell-blocks of Strain 1 (above) and Strain 2 (below) dividing towards (shown in color) and away from (shown in black) cell-blocks of the other strain with the threshold AHL amount to induce gene expression in Strain 1 being varied and the threshold for Strain 2 kept constant at 5 nM.

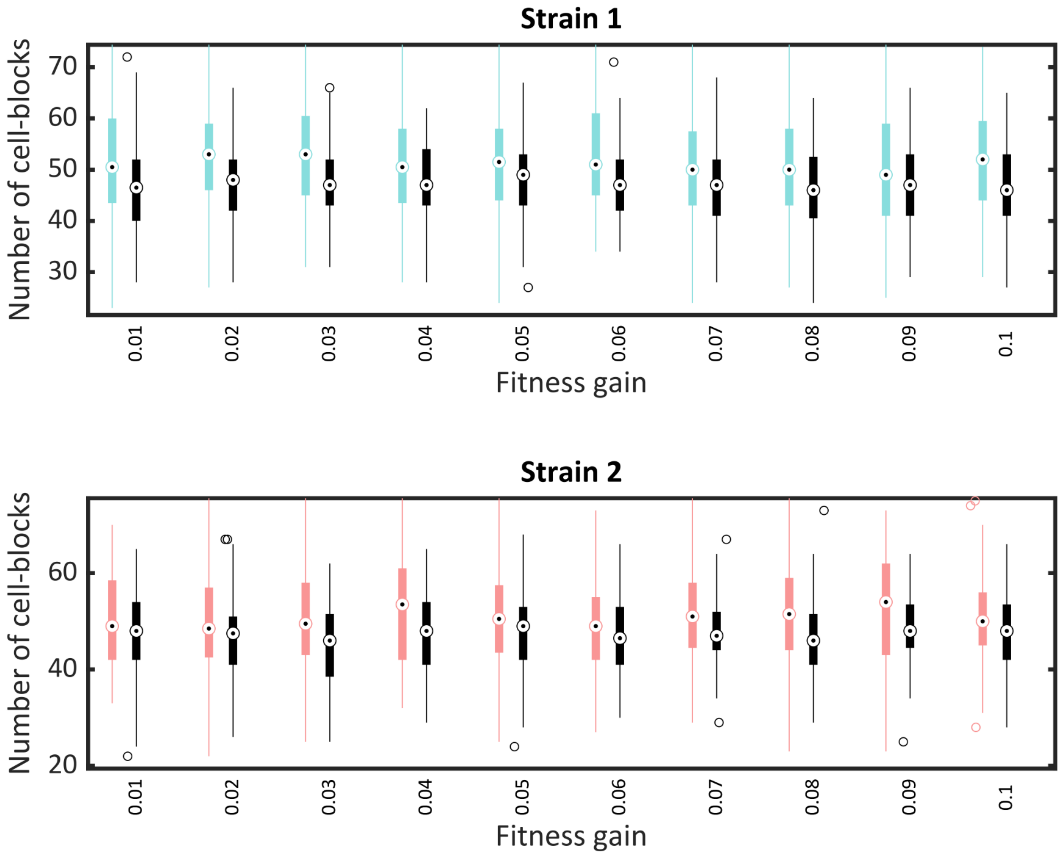


**Supp. Fig. 2**: Results of a preferential growth assay quantifying the number of cell-blocks of Strain 1 (above) and Strain 2 (below) dividing towards (shown in color) and away from (shown in black) cell-blocks of the other strain with the fitness gain due to QS-mediated gene expression in Strain 1 being varied and the fitness gain for Strain 2 kept constant at 0.05.

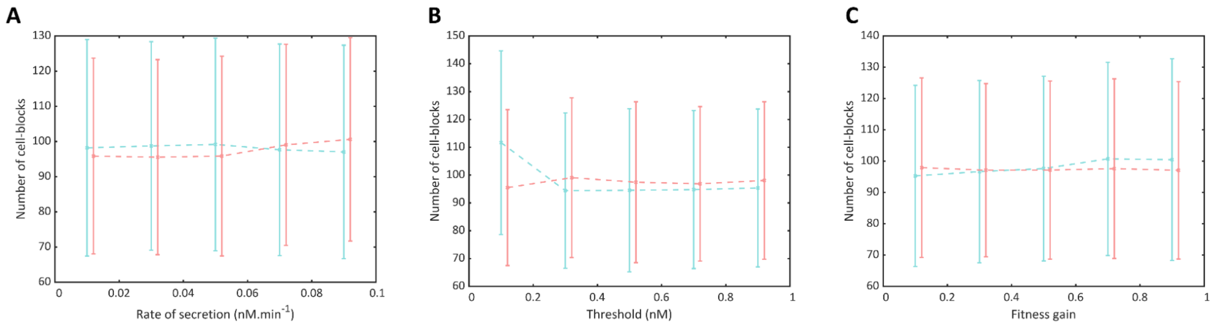


**Supp. Fig. 3**: Results of the parameter grid search. (A) The populations of each strain in the two-strain QSMM with the rate of secretion of AHL molecules from Strain 1 being varied and the rate of secretion from Strain 2 kept constant at $1.6\times10^{-2}$ nM cell-block$^{-1}$ min$^{-1}$. (B) The populations of each strain in the two-strain QSMM with the threshold AHL amount to induce gene expression in Strain 1 being varied and the threshold for Strain 2 kept constant at 5 nM. (C) The populations of each strain in the two-strain QSMM with the fitness gain due to QS-mediated gene expression in Strain 1 being varied and the fitness gain for Strain 2 kept constant at 0.05.

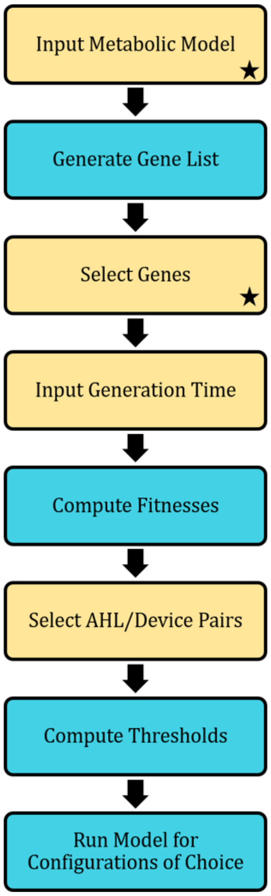


**Supp. Fig. 4**: Flowchart depicting the workflow of our models in the package. Steps in yellow indicate that user input is required while steps in blue indicate the background computations. The starred steps indicate the points from which the workflow can be initiated.
